# Influence of social and semantic contexts on phonetic encoding in naturalistic conversations

**DOI:** 10.1101/2024.01.10.575068

**Authors:** Etienne Abassi, Robert J. Zatorre

**Affiliations:** Montreal Neurological Institute – McGill University, Centre for Research in Brain, Language and Music (CRBLM) International Laboratory for Brain, Music and Sound Research (BRAMS)

**Keywords:** Phonetic encoding, Conversations, Context, Social, Semantic

## Abstract

Social interactions occupy a significant part of life, and understanding others’ conversations is key to navigating our social world. While the role of semantics in speech comprehension is well-established at the word or sentence level, its influence on larger conversational scales, alongside social context, is less understood. The present study examined how semantic and social contexts modulate phonetic encoding during natural conversations using a speech-in-noise paradigm. Participants listened to AI-generated dialogues (two speakers) or monologues (one speaker) in an intact or sentence-scrambled order. Each trial contained five sentences, with the fifth sentence embedded in multi-talker babble noise. The same sentence was then repeated without noise, with one word either altered or unchanged. Healthy adults identified whether the sentence matched the in-noise version. Through several online experiments (N=211), both social and semantic contexts showed influences on speech-in-noise processing, with improved performance for dialogues over monologues and for intact over sentence-scrambled conversations. These results suggest that both semantic and social factors shape speech comprehension, emphasizing their role in auditory cognition. This finding raises important questions about predictive and other mechanisms involved in processing complex, multi-sentence conversations, underscoring the critical role of social interaction in communication.

## I. INTRODUCTION

During our lifetime, we humans deploy complex abilities to navigate our social world (Adolphs, 2001, 2009). Consequently, a significant portion of our existence is devoted to engaging in social interactions with others. Not only do we interact in the first person, but listening to others’ interactions also serves critical functions in understanding our social world. Hence, successful understanding of others relies on the seamless integration of sensory input and cognitive mechanisms, enabling us to understand speech across diverse and often challenging auditory environments. Central to this process is phonetic encoding, the mapping of acoustic signals onto linguistic representations, which underpins speech comprehension (Clements, 1985; Jakobson et al., 1951; Lahiri & Reetz, 2010; Sapir, 1925; Yi et al., 2019). While much of the existing research has examined how speech is processed in controlled environments using single sentences or words, real-world communication involves multiple layers of complexity, including listening contexts that influence how phonetic encoding unfolds (Kuperberg & Jaeger, 2016; Sohoglu et al., 2012). Understanding the mechanisms by which these contexts shape speech perception is critical for advancing theories of auditory cognition and improving interventions for individuals with hearing loss.

Semantic context, or the coherence and meaning embedded in speech, has long been recognized as a critical factor in speech processing, and plays a pivotal role in phonetic encoding. Coherent sentences allow listeners to predict upcoming words and phonetic structures, reducing cognitive load and improving processing efficiency (Sohoglu et al., 2012; Zekveld et al., 2011, 2012, 2013). For example, sentences with meaningful content enable listeners to disambiguate phonetic details, particularly in acoustically challenging environments such as noisy or multi-talker settings (Kuperberg & Jaeger, 2016; O’Neill et al., 2021). These predictive mechanisms are often conceptualized within the framework of predictive coding, which posits that the brain generates expectations based on prior input to minimize the mismatch between sensory input and cognitive predictions (Clark, 2013; Friston, 2005; Rao & Ballard, 1999). Semantic congruence facilitates phonetic encoding by reinforcing expectations and enhancing signal clarity, while semantic incongruence or nonsense sentences disrupt these processes and impose greater cognitive demands (Golestani et al., 2009, 2013; Zekveld et al., 2011).

Research on semantic context has typically focused on isolated words or sentences. For instance, studies have shown that listeners process phonemes more accurately when words appear in semantically predictable sentences (Roland et al., 2012; Zekveld et al., 2013) and that speech patterns at different timescales are processed using different mechanisms and/or brain circuitry. Indeed, shorter timescales, such as phonemes and syllables, primarily engage the left middle superior temporal gyrus (mid-STG), while longer units like words or sentences recruit the left anterior STG (DeWitt & Rauschecker, 2012; Lerner et al., 2011). Additionally, EEG recording during speech naturalistic listening shows that semantic contexts can enhance early auditory responses as well as speech envelope reconstruction (Broderick et al., 2019). However, most work to date has examined semantic processing in highly constrained contexts, such as isolated words, word pairs, or single sentences presented in controlled, low-variability environments. While these approaches offer experimental precision, they do not fully capture the richness of everyday language use. In natural speech, particularly in conversation, meaning is built incrementally across multiple utterances, supported by broader semantic and pragmatic cues. Studies using connected speech materials, such as the Connected Speech Test (Cox et al., 1987), a standardized measure of speech intelligibility using short conversational passages presented in background noise, or naturalistic corpora like the LUCID database (Baker & Hazan, 2010, 2011), suggest that broader semantic context improves intelligibility. However, these studies typically do not isolate the specific effects of context on phonetic encoding. The present study addresses this gap by examining how semantic context within prototypical conversations influences phonetic encoding. Here, we focus on how large-scale semantic congruency in the context of conversations can support lower-level auditory encoding, shedding light on the interaction between linguistic prediction and speech perception in more ecologically valid settings.

While semantic context focuses on the linguistic coherence of speech, social context, on the other hand, encompasses the interactional dynamics and relational cues present in communication. As such, it may represent another critical factor influencing phonetic encoding. Here, dynamic exchanges between speakers, such as turn-taking and conversational flow, may engage cognitive systems that are tuned to socially relevant stimuli while providing listeners with additional scaffolding for auditory processing. For instance, dialogues involving alternating speakers provide structural and temporal cues that enhance speech comprehension compared to monologues (Fox Tree, 1999). These interactional dynamics may facilitate phonetic encoding by directing attention to relevant acoustic signals and reinforcing cognitive predictions about speaker behavior (Levy et al., 2003). Social context is further enriched by factors such as speaker identity, familiarity, and social relationships. Familiar voices are processed more efficiently than unfamiliar ones (Domingo et al., 2020; Johnsrude et al., 2013), as listeners can leverage prior knowledge to disambiguate phonetic details and interpret speech more effectively (Belin et al., 2000). Similarly, shared social knowledge and relational dynamics can modulate auditory processing by providing additional contextual scaffolding. For example, listeners process speech more accurately when they share a common linguistic background with the speaker (Adank et al., 2009; Lev-Ari & Keysar, 2010; Smith & Bisazza, 1982), although the overall evidence for interlanguage benefits is mixed, with several studies reporting only marginal or null effects (Bent & Bradlow, 2003; Hayes- Harb et al., 2008; Major et al., 2002; Munro et al., 2006). On a last note, social context can also influence speech perception not only through interactional dynamics, but also via social evaluation. Indeed, prior work has shown that listeners’ perceptions of intelligibility are shaped by attitudes, stereotypes, and expectations about the speaker, even when acoustic cues are held constant (Babel & Russell, 2015; Lev-Ari & Keysar, 2010; McGowan, 2015; Niedzielski, 1999; Rubin, 1992). These findings highlight the broader role of social cognition in auditory processing (Drager & Kirtley, 2016). Overall, these effects suggest that social interaction engages specialized neural and cognitive systems that enhance phonetic encoding and speech comprehension. Dialogues, as a prototypical form of social interaction, offer particularly rich opportunities for examining the influence of social context. Unlike monologues, dialogues involve interactive turn- taking, which creates temporal predictability and facilitates attentional engagement. Previous studies have shown that listeners perceive dialogues as more intelligible than monologues, even when the content is identical, suggesting that interactional dynamics enhance cognitive processing (Branigan et al., 2011; Pickering & Garrod, 2021; Tolins et al., 2018). However, the mechanisms underlying this advantage remain poorly understood.

Going further, the influence of context in sensory processing of a stimulus is not only known in the auditory modality but also in the visual modality (Brandman & Peelen, 2017; Oliva & Torralba, 2007), where recent work showed that the neural representation of a single body within a dyad is sharpened when the two bodies are represented face-to-face (as if interacting) as compared to back-to-back (Abassi et al., 2020; Bellot et al., 2021). This effect shows that a given social context can have a significant influence on the perception of a visual stimulus and its neural processing. Both visual and auditory systems are organized hierarchically, progressing from early sensory encoding (e.g., pitch or shape) to the integration of higher-level features such as linguistic content or social intent (Iamshchinina et al., 2022). These layered structures allow top-down influences, such as contextual or social expectations, to shape perception in both modalities. For instance, just as the presence of an interacting body pair enhances neural encoding of a single body in vision (Abassi et al., 2020), we hypothesize that dialogue-based interaction may enhance encoding of verbal sentences in auditory scenes. Drawing on this analogy, we suggest that social context may similarly modulate auditory perception, particularly under challenging listening conditions. Here, the social context involves intricate mechanisms related to the interplay between two or more speakers, elements that are generally overlooked in the examination of higher-order top-down cues. Thus, building on the literature of both social audition and vision, a major aim of this study is isolating the effects of social interaction on phonetic encoding, controlling for semantic content to focus on the contribution of interactional cues such as speaker alternation and conversational flow.

On a last point, individual differences in a listener’s sensitivity to social contexts may influence phonetic encoding, particularly through variations in social traits. Indeed, the Autism Spectrum Quotient (AQ), a widely used measure of autism-related traits in both neurodiverse and neurotypical populations (Baron-Cohen et al., 2001), has been consistently linked to reduced sensitivity to social stimuli. More specifically, neurotypical individuals with higher AQ scores demonstrate diminished sensitivity in visual tasks involving social stimuli, such as processing dyadic body interactions (Abassi & Papeo, 2022), faces (Wyer et al., 2012), biological motion (van Boxtel et al., 2017), and animals (Yang et al., 2024). These findings suggest a general link between individual differences in social-related traits and integration of social cues in the visual modality, which may extend to auditory perception in contexts emphasizing social interaction dynamics. In parallel, higher AQ scores have been associated with enhanced attention to fine phonetic details (Ota et al., 2015; Stewart & Ota, 2008; Yu, 2010), suggesting a potential advantage in auditory tasks focusing on precise acoustic distinctions. However, this strength may not fully compensate for the reduced sensitivity to social dynamics in tasks requiring the integration of social cues for phonetic processing. This dual influence highlights how individuals with higher AQ scores might excel in isolated phonetic tasks but face challenges in contexts where social dynamics are critical for guiding perceptual processes. By including AQ scores in the present study, we aim to investigate how sensitivity to social interaction dynamics may modulate the effects of semantic and social contexts on phonetic encoding.

Overall, in the current study we investigated the influence of semantic and social contexts on phonetic encoding. Specifically, we hypothesized that both semantic coherence and dynamic social interactions would enhance the encoding of phonetic information. In addition, we also sought to explore whether their effects are independent or interacting. To test these hypotheses, we manipulated the semantic context (intact vs. sentence-scrambled conversations) and social context (dialogues vs. monologues) in which a sentence was heard, using a speech-in-noise paradigm to measure perceptual sensitivity. Speech-in-noise paradigms provide an effective framework for studying phonetic encoding under controlled conditions that approximate real-world listening challenges. By introducing competing auditory stimuli, these tasks require listeners to draw on higher-order cognitive mechanisms, such as context-based predictions, to process speech effectively (Zekveld et al., 2013). In our design, semantic coherence may primarily enhance phonetic encoding by increasing linguistic predictability, whereas social context may offer cues related to speaker dynamics and turn-taking structure. While our focus is on identifying the contribution of each context, our factorial design also allows us to test whether their effects are independent or interacting. By combining speech-in-noise tasks with controlled manipulations of semantic and social cues, we aimed to isolate the mechanisms through which large-scale high-level contexts could enhance speech perception. However, unlike previous research, which has often focused on isolated words or sentences, our study examines how these factors operate at the level of larger-scale naturalistic conversations, offering a more comprehensive understanding of real-world speech perception.

To address these questions, we conducted two main experiments (summarized in Table 1). Experiment 1 was a preliminary study aimed at identifying an appropriate signal-to-noise ratio (SNR) for use in the main experiment. Experiment 2 tested the effects of semantic and social context on speech processing across several presentation formats, using different groupings and block structures to control for potential order effects. The implications of this research extend beyond basic auditory science, as individuals with hearing loss or social-cognitive impairments often struggle to process speech in complex environments where both semantic and social cues play a critical role (Ciorba et al., 2012; Nordvik et al., 2018). By elucidating the mechanisms underlying phonetic encoding in these contexts, this study lays the groundwork for developing more ecologically valid assessment tools and assistive technologies, such as hearing aids that leverage conversational dynamics and contextual predictions to enhance speech processing.

**Table I.**
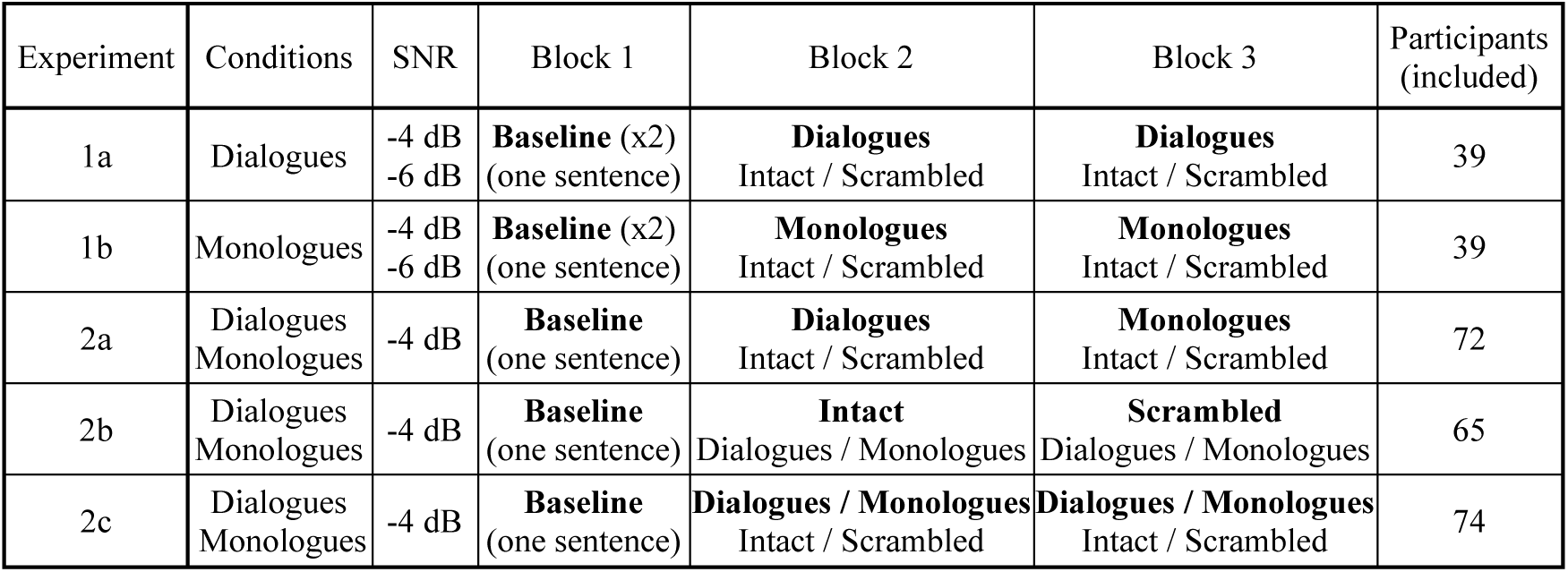
Summary of specificities of designs, conditions and signal-to-noise ratios (SNRs) for each experiment.

| Experiment | Conditions | SNR | Block 1 | Block 2 | Block 3 | Participants (included) |
| --- | --- | --- | --- | --- | --- | --- |
| 1a | Dialogues | -4 dB<br>-6 dB | **Baseline** (x2)<br>(one sentence) | **Dialogues**<br>Intact / Scrambled | **Dialogues**<br>Intact / Scrambled | 39 |
| 1b | Monologues | -4 dB<br>-6 dB | **Baseline** (x2)<br>(one sentence) | **Monologues**<br>Intact / Scrambled | **Monologues**<br>Intact / Scrambled | 39 |
| 2a | Dialogues<br>Monologues | -4 dB | **Baseline**<br>(one sentence) | **Dialogues**<br>Intact / Scrambled | **Monologues**<br>Intact / Scrambled | 72 |
| 2b | Dialogues<br>Monologues | -4 dB | **Baseline**<br>(one sentence) | **Intact**<br>Dialogues / Monologues | **Scrambled**<br>Dialogues / Monologues | 65 |
| 2c | Dialogues<br>Monologues | -4 dB | **Baseline**<br>(one sentence) | **Dialogues / Monologues**<br>Intact / Scrambled | **Dialogues / Monologues**<br>Intact / Scrambled | 74 |

## II. METHODS

### A. Participants

A total of 295 participants took part in our on-line study (Exp 1: 80 participants, 29 females, 3 sex unreported, mean age 29.8 years +/- 6.2 SD; Exp 2: 215 participants, 85 females, 2 sex unreported, mean age 31.6 years +/- 5.6 SD). Overall, the minimal sample size for each experiment was based on previous literature using analogous word-recognition tasks in multi-talker babble noise. More specifically, Killion et al. (2004) found significant results for the QuickSIN test (which measures whole sentences recognition in noise) using 16 participants and Wilson (2003) used 24 participants for the Word-In-Noise (WIN) test (which assesses isolated words recognition in noise), while in our current study we had a minimum of 40 participants for each experimental subset (see below). All participants read and signed an informed consent form prior to participating in the study and were remunerated 6£ for ∼40 minutes of their time, paid through the on-line platform (prolific.com). They were all American English first-language speakers with normal hearing (determined by self-report online), aged between 18 and 40 years. They were recruited on the Prolific platform and the study was restricted only to people with a desktop computer or a laptop. Prior to enrolling in the study, online participants were instructed that wearing headphones or earbuds was mandatory to participate. They were also asked to be in a silent and calm environment, as well as avoid having any object around them that may disturb their attention, such as a smartphone or a TV on.

### B. Stimuli

#### 1. Stimuli features

The main stimulus set consisted of AI generated English two-speaker dialogues or one-speaker monologues (factor: social context) arranged in intact or sentence-scrambled order (factor: semantic context) (Fig. 1). In the current study, we use the term “conversation” as a general label to describe all four stimulus conditions, for the sake of simplicity and consistency across the manuscript. However, only the intact dialogue condition fully meets the conventional definition of a “conversation”, featuring multiple speakers, turn-taking, and coherent topical flow. The other conditions systematically altered aspects of this structure: In the monologue conditions, the same dialogues were rendered by a single speaker, thus preserving sentence content but eliminating speaker alternation. In the scrambled conditions, sentence order was randomized, disrupting semantic coherence at the conversational level while maintaining semantic coherence at the sentence-level. All stimuli were created using a combination of ChatGPT (GPT-3.5, *OpenAI*) to generate the contents of the conversations and Google text-to-speech AI to convert the textual conversations into natural-sounding speech. The entire set of stimuli and materials for generation can be found online at www.zlab.mcgill.ca/downloads/Stimuli_from_Abassi_2025. A sample is also provided as supplementary files (*SuppPub2.zip*).

#### 2. Speech voices AI generation

The choice of using AI-generated stimuli instead of recorded human actors voices was made for two main reasons: 1) AI generation allowed us to create auditory stimuli that were purely prototypical of real speech and, importantly, devoid of any fluctuation in the inner features of speech that naturally occurred from humans’ speakers (i.e., variations in rhythm, intonation, articulation, etc…), that could have created confounds between sentences or conditions. This point was especially important in light of the specific conditions we were studying, for which the individual sentences features had to be exactly identical across conditions. 2) AI generation allowed us to have very fine control of specific features of speech that we wanted to be able to easily modulate (i.e., fundamental frequency, pitch and prosody). AI-generated speech may lack features of truly natural conversation, such as prosodic entrainment across speakers or adaptive rhythm over time. However, our goal was not to simulate fully natural conversations, but to create prototypical and tightly controlled conversational stimuli, especially with regard to timing, pitch range, and segmental clarity. For this reason, we prioritized consistency and acoustic control over naturalness *per se*, and we did not conduct formal perceptual validation. Informal pilot testing confirmed that the speech was easily understood by naïve listeners and that it sounded quite natural.

**Figure 1.**
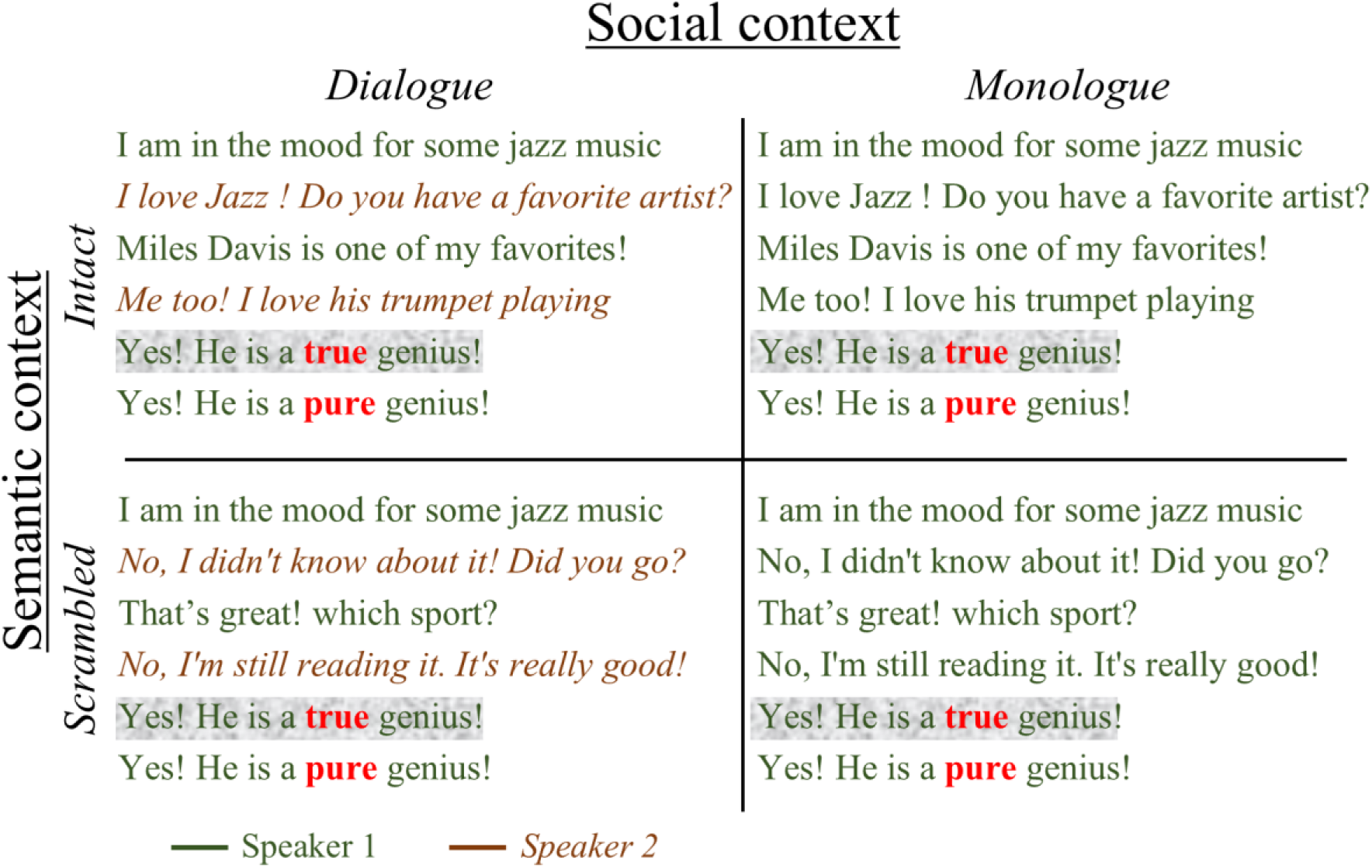
Example of conversations composing the stimuli set. Stimuli are arranged following a 2×2 design with factors social context (Dialogue / Monologue) and semantic context (Intact / Scrambled). For dialogues, sentences presented in green standard text alternating with brown italic text represent the alternation between the two speakers (one female and one male). Sentences presented over grey background illustrate that the fifth sentence of each conversation was always embedded in noise. The last sentence could be either the same or different than the previous one, with only one word changed between the sentences, as represented in bold red. The entire set of stimuli can be found online at www.zlab.mcgill.ca/stimuli (‘Stimuli_from_Abassi_2025’) or attached in supplementary materials (*SuppPub2.zip*).

We selected a fundamental frequency (f0) of 120 Hz for the male voice, and of 181 Hz for the female voice using the built-in-function of Google text-to-speech AI that permits to increment or decrement the overall pitch of a voice by a custom number of semi-tones. Thus, for a chosen f0, we looped between the text-to-speech AI generator and custom MATLAB codes to measure the f0, by decrementing or incrementing by one semitons until we reached our chosen f0. Here, the chosen f0 values for the female and male voices were separated by 4 semi-tones, and this two f0 were chosen in a set of 10 semi-tones ranging from 120 Hz to 203 Hz. We selected this overall f0 range so it was large enough to cover a representative range of f0 in the general population (see Cristina Oliveira et al., 2021). The remaining semi-tones were used as f0 for the 8 competing voices used as multi-talker babble noise, for which details are given below (see also supplementary figure 1, featuring a graphical similarity representation of the distribution of semi-tones across male and female voices). For the voice’s identities used in the final generation, we chose 2 voices among a list of 10 proposed voices by the text-to-speech generator (male voice identity: “en-US- Wavenet-J”; female voice identity: “en-US-Wavenet-G”), with the criteria of their initial f0 before fine-tuning being the closest to the f0 we aimed to reach.

#### 3. Speech content AI generation

Each conversation was then carefully constructed to ensure that dialogues and monologues had the same sentence structures and themes, isolating the effect of social context. Indeed, dialogues featured alternating male and female voices, providing turn-taking cues and interactive dynamics that were absent in monologues, which were all produced by the same voice. This design allowed us to investigate how the presence of two interacting voices, versus a single continuous voice, influences the listener’s processing of speech. More specifically, we first used ChatGPT to generate the text of 35 five-sentence conversations based on five themes (Music, Cinema, Sport, Literature and Travel). An example of the ChatGPT prompts used is *“On the topic of music, write X conversations of 5 sentences each between two persons with a maximum of 15 words per sentence”*. We did not use the exact conversation contents as generated by ChatGPT, but rather made modifications such as word, syntax, and length changes to enhance the naturalistic aspect of each item. For each five-sentence conversation, we manually created a sixth sentence by changing a single word from the original (fifth) sentence. The change was designed to be subtle, maintaining grammaticality and plausibility within the sentence while introducing a distinct phonetic form. For example, *“He is a **true** genius”* was changed to *“He is a **pure** genius”* The substituted words were chosen to differ phonologically from the original, while avoiding semantic or pragmatic violations. To avoid predictability, the location of the altered word varied across sentences (beginning, middle, or end) and was not fixed. The modified words had the same number of syllables as the target word and were drawn from the same broad grammatical category (typically adjectives, nouns, or verbs). This semi-controlled approach allowed for natural-sounding stimuli while minimizing systematic cues. Thus, the manipulations were exclusively related to the *context* within which the target sentence was presented, and not to the structure or content of the target sentence itself.

#### 4. Prosody generation

To enhance prosodic naturalness, we used Speech Synthesis Markup Language (SSML), an XML-based standard developed by the World Wide Web Consortium (W3C) to control expressive features of synthetic speech, such as pitch, speaking rate, volume, and pausing (W3C, 2010). SSML allows users to insert markup tags into text that guide how speech synthesis engines render prosody. In our case, we used it to manually adjust pitch and rate within sentences. For example, we used *<prosody rate=“fast” pitch=“+4st”>* to increase speed and pitch on specific words. While SSML does not encode linguistic prosodic features (like pitch accents or boundary tones) directly, it provides precise control over acoustic cues. This helped ensure that individual sentences were prosodically varied and easily interpretable, even if the resulting speech did not fully replicate the dynamic prosody of natural human dialogue. The complete list of sentences with their prosody tags is available in the stimuli dataset package given in supplementary materials (*SuppPub2.zip*). We then selected two voices from the list proposed by Google text-to-speech AI: one female voice (*‘en-US-Wavenet-G’*) and one male voice (*‘en-US-Wavenet-J’*). We finally converted each text sentence of each conversation into auditory speech for the two voices, and we normalized all audio tracks to have the same root mean squared (RMS) amplitude across all sentences.

#### 5. Sentences arrangements

We then arranged the sentences into conversations as follows. For intact dialogues, half of the conversations began with the female voice and alternated with the male voice, and for the other half it was the opposite. For monologues, half of the conversations were composed of the female voice only, while the other half was composed of the male voice only. Furthermore, for both dialogues and monologues, sentence-scrambled conversations were created by randomly shifting sentences across the conversations (i.e., across distinct topics), except for the fifth sentences. Sentence positions were also fully randomized within each conversation, rather than swapped across conversations, to prevent accidental preservation of overarching structures such as question-answer pairs or turn-taking patterns.

In total, we created 240 conversational stimuli (60 stimuli by unique condition, half primed with female voice and half primed with male voice). Between every sentence, we added a silent gap randomized between 400 and 600 ms, except between the fifth and the sixth sentences, where a silent gap of 1 s. was added. On average, each full conversation had a total duration of ∼18 s.

#### 6. One-sentence stimuli

In addition to dialogues and monologues, we created a set of 50 one-sentence stimuli that were different than the sentences used in the conversations and were used as a baseline condition to control for inter-individual variability in performance in processing speech-in-noise. For each of these sentences, we created a second sentence by hand, that was the same as the first sentence with the change of a single word from the original sentence, and the location of the altered word varied across sentences (beginning, middle, or end). Similarly to the main experiment (conversations), the word change itself was unrelated to the content of the conversation—that is, it was a neutral word within the sentence that would be equally appropriate in the context. We then converted these sentences into natural-sounding speech for both female and male voices, using the same method as for the conversations. For example, one of the ChatGPT prompts used to generate the text of these one-sentence stimuli was *“Write X sentences on the topic of music with a maximum of 15 words per sentence”.* We then added prosody tags following SSML rules and transformed them into audio using Google text-to-speech. In total, we created 100 one-sentence stimuli (50 for female and 50 for male), for an average duration of ∼5 s. each.

#### 7. Multi-talker babble noise

In a last step, we embedded the fifth sentence of each conversation, as well as all the one-sentence baseline stimuli into 8-voices multi-talker babble noise. We used multi-talker babble noise as a masker for our stimuli because we wanted it to be as naturalistic as possible (for a review on the different types of maskers, see Assmann & Summerfield, 2004)). Detailed information about the noise features and its generation are given in supplementary information 1. After generating the 24 audio distractor sentences for each of the 8 voices, we created the multi-talker noise stimuli using custom Matlab codes, closely following the method of Van Engen & Bradlow (2007): 1) Each audio track was normalized to have the same RMS amplitude across all tracks; 2) For each talker, two randomly chosen sentences were concatenated to ensure the duration of the noise tracks would exceed the durations of all target sentences; 3) A random multiple of 100 ms (0–500 ms) of silence was added at the beginning of each concatenated sentences to stagger the talkers once they were mixed together; 4) All eight talkers were mixed, by randomly choosing one concatenated sentence by talker, and the initial 500 ms of the concatenated sentences was removed to eliminate noise that did not contain all eight talkers; 5)

The first 100 ms of the complete noise tracks was faded in. In total, we produced 120 multi-talker noise stimuli. In a final step, we mixed the fifth target sentences of each of the previously generated conversations or each of the one-sentence stimuli with a randomly chosen noise stimulus for the entire duration of the target sentences with the addition of 100ms before and after the sentences during which there was only noise. Here, the level of the multi-talker masking noise was held constant, while the amplitude of the target stimuli was decreased by Signal-to-Noise Ratios (SNRs) of -1, -4, and −6 dB, where -4 and -6 dB are known to produce a speech reception threshold (see Plomp & Mimpen, 1979) that is in the range of 40 % to 60 % depending on the studies and methods (Brungart et al., 2001; Dingemanse & Goedegebure, 2020; Festen & Plomp, 1990; Hagerman, 1984; McArdle et al., 2005; Miller et al., 1951). These values were also tested in prior in-the-lab pilot testing (unpublished data) to estimate a range of SNRs that would adequately capture the skill level of young healthy participants, while limiting floor or ceiling effects as well as subject frustration (Coffey et al., 2019).

## D. Procedure

### 1. Overview

We used a speech-in-noise paradigm to study how semantic and social contexts interact with phonetic encoding of a sentence. Importantly, we used a task that required the listener to identify if a word had been changed, but that word was not itself relevant to the content of the dialogue (e.g., see figure 1, in which the changed word was “true” vs “pure”, both of which fit the meaning of the sentence, and neither of which is related to the dialogue’s content). This manipulation allowed us to focus on the phonetic encoding of a word rather than its underlying meaning, moving beyond traditional measures of intelligibility by focusing on the deeper cognitive processes that enable robust speech perception. Here, we note that we didn’t explicitly ask the participants to perform the task on a specific word change, but rather to identify a change between the original sentence and the repeated one. The exact instruction given to the participant was *“Your task is to determine, as quickly as possible, whether the repeated sentence is exactly the same or different than the one presented in noise. It’s crucial to note that even subtle changes between the repeated sentences should be reported as different. Only when two sentences are precisely identical should you consider them the same”*.

### 2. Experiments rationales

We conducted two main experiments (see table I for a summary of the experiments and their specificity). Experiment 1 was a preliminary study designed to select a correct SNR (-4 dB or -6 dB) to be used in the subsequent experiments and was composed of two participant groups: one with dialogues only (Exp. 1a) and one with monologues only (Exp. 1b). Experiment 2 addressed the effects of semantic and social contexts while controlling for effects of different experimental conditions/orders (Carvalho & Goldstone, 2014a, 2014b; Francis & Ciocca, 2003), and was composed of 3 participant groups: Experiment 2a was composed of separate blocks of dialogues-only trials and blocks of monologues-only trials, each featuring both intact and sentence-scrambled conversations within each block; Experiment 2b was composed of separate blocks of intact conversations only and blocks of sentence-scrambled conversations only, each featuring both dialogues and monologues within each block; and Experiment 2c was composed of blocks featuring all four conditions within each block. All experiments (for which details are given below) were developed using PsychoPy (Peirce et al., 2019), then transformed into JavaScript using the PsychoJS library as implemented in PsychoPy and hosted on the Pavlovia platform (www.pavlovia.org) for online running of the experiments.

### 3. Experimental procedure

Prior to beginning an experiment, participants went through a series of checks to ensure optimal hearing conditions while performing the tasks (Zhao et al., 2022), for which details are given in supplementary information 2. Following this, participants had to read and sign electronically the consent form, after which they were allowed to participate in the study. Each experiment (for which specific differences are given below) was preceded by a training task that was exactly the same as the testing task. For each training task, sentence-in-noise SNRs were of -1 dB for the first half of the stimuli and of -4 dB for the second half, while each training trial was followed by feedback. Each experiment consisted of two main tasks: 1) a baseline task, in which only one-sentence stimulus was presented in noise, that was used to control the level of understanding of sentences-in-noise for each participant without prior social or semantic context given, and 2) a conversational task, in which dialogues or monologues in intact or sentence-scrambled order were presented. For both tasks, immediately after hearing a sentence in noise, the participants heard it again but without noise. Half of the time, these repeated sentences were exactly the same as the one presented with noise, while the other half had a one-word difference. The task was to determine, as quickly as possible, whether the repeated sentence was exactly the same or different than the one presented in noise. A typical trial began with a waiting time of 500 *ms*. with a fixation cross at the center of the screen, followed by the presentation of the auditory stimulus, then by a visual reminder of the response buttons until the participants pressed one of the two buttons. Every 32 trials, the participants were invited to take a break. The response buttons were counterbalanced across participants (i.e., half of the participants used the left arrow for the ‘same’ trials and right arrow for the ‘different’ trials, while the other half did the opposite). The first voice heard was also counterbalanced across participants (i.e., half heard the female voice first in the first presented stimuli, while the other half heard the male voice in first). For all tasks, the order of trials was pseudo-randomized to avoid the presentation of more than 3 of the same trial type (same/different), target voice (male/female), social factor (dialogue/monologue), semantic factor (intact/scrambled) or SNR (-4/-6 dB for Exp. 1 only). At the end of the experiment, participants completed the 50-item Autism-Spectrum Quotient scale (AQ; Baron-Cohen et al., 2001), for which details and rationale are given below.

All responses (Accuracy, reaction time [RTs] and AQ questionnaire) were collected with Pavlovia and processed with Matlab. Measure of reaction times began directly after the last sentence was heard until the participant pressed one of the two response buttons.

### 4. Experimental designs

#### Experiment 1

Experiment 1 was conducted to choose a correct SNR for measuring our effects of interest in the subsequent experiments and included two groups, with one group presented with dialogues only (Exp. 1a; 40 participants) and one group presented with monologues only (Exp. 1b; 40 participants). Here, we expected a correct SNR to not produce ceiling of floor effects with accuracy scores and to produce our effect of interest (i.e. better performance for intact over sentence-scrambled conversations using *d’* scores and/or RTs). For each group, half of the unique stimuli were presented with an SNR of -6dB, while the other half was presented with an SNR of -4dB, and stimuli were randomly assigned in one or the other half for each participant. The alternation of -4dB and -6dB stimuli was randomized. For the two groups, the experiment began with two training tasks: one with 8 baseline trials and the other with 4 conversation trials. The order of the training tasks was counterbalanced over participants. Following the training, participants underwent the experiment that consisted of two baseline blocks (32 trials each) and two conversation blocks (32 trials each) interleaved, for which the order was counterbalanced across participants.

#### Experiment 2a

Experiment 2a included 74 participants and was composed of blocks of dialogues only or monologues only, while a same block featured both intact and scrambled conversations. For each participant, half of the unique stimuli was presented in the dialogue condition and repeated across the intact and scrambled conditions while the other half was presented in the monologue condition and repeated across the intact and scrambled conditions, and stimuli were randomly assigned in one or the other half for each participant using 6 predefined lists. Experiment 2a used an SNR of -4 dB and was conducted as follows. At first a training task featured 6 baseline trials, followed by a testing task of 44 baseline trials. Afterwards, participants were presented with one block of dialogues only and one block of monologues only (32 trials each, with half intact and half sentence-scrambled conversations, presented in random order). The order of blocks was counterbalanced across participants. Each block was preceded by a training task of 4 trials of the same condition.

#### Experiment 2b

Experiment 2b included 66 participants and was composed of blocks of intact conversations only or sentence-scrambled conversations only, while a same block featured both dialogues and monologues. For each participant, half of the unique stimuli was presented in the intact condition and repeated across dialogues and monologues while the other half was presented in the scrambled condition and repeated across dialogues and monologues, and stimuli were randomly assigned in one or the other half for each participant using 6 predefined lists. Experiment 2b used an SNR of -4 dB and was conducted as follows. At first a training task featured 6 baseline trials, followed by a testing task of 44 baseline trials. Afterwards, participants were presented with one block of intact conversations only and one block of sentence-scrambled conversation only (32 trials each, with half dialogues and half monologues presented in random order). The order of blocks was counterbalanced across participants. Each block was preceded by a training task of 4 trials of the same condition.

#### Experiment 2c

Experiment 2c included 75 participants and was composed of blocks featuring all conditions randomly intermixed (i.e., both dialogues and monologues in both intact and sentence-scrambled orders, not blocked into conditions). For each participant, half of the unique stimuli were presented in the dialogue condition and repeated across the intact and scrambled conditions while the other half were presented in the monologue condition and repeated across the intact and scrambled conditions, and stimuli were randomly assigned in one or the other half for each participant using 6 predefined lists. Experiment 2c used an SNR of -4 dB and was conducted as follows. At first a training task featured 6 baseline trials, followed by a testing task of 44 baseline trials. Afterwards, participants were presented with two blocks (32 trials each) featuring all conditions.

### 5. Autism Quotient (AQ) task

For experiment 2, at the end of each acquisition participants completed the 50-item Autism-Spectrum Quotient scale (AQ; Baron-Cohen et al., 2001) which measures the degree to which an adult with normal intelligence shows traits associated with the ASD (Autistic Spectrum Disorders), in the domain of social skills, attention-switching, attention to details, communication, and imagination. Subjects responded to each item using a 4-point rating scale, ranging from “definitely agree” to “definitely disagree”. Calculation of the AQ quotient was performed following the instructions provided by Baron-Cohen et al. (2001). Higher scores indicate greater ASD traits while lower scores indicate fewer ASD traits.

## E. Data preprocessing

For all experiments, the following data preprocessing steps were applied: 1) For each participant, trials for which RTs were >3 SD above the participant mean were discarded from further analyses; 2) For each participant, trials in which the response was inaccurate were discarded for RTs only; 3) Participants were excluded from the study if their mean RTs averaged across all conversation conditions was >3 SD above the subgroup mean (Participants exclusions per experiment: *Exp. 1a:* 1/40; *Exp. 1b:* 1/40; *Exp. 2a:* 2/74; *Exp. 2b:* 1/66; *Exp. 2c:* 1/75; *Total of participants exclusions:* 6/295; ∼2.03 %). After exclusions, statistical analyses for all conditions within an experiment included 39 participants for experiment 1a, 39 participants for experiment 1b and 211 participants for experiment 2.

## F. Analyses

### 1. Main analyses

For all experiments, d-prime scores (*d’*) and mean RTs were used to compare the effect of intact vs. scrambled conversation across dialogues and monologues. Here, we computed *d′* scores for each participant as this measure, derived from signal detection theory, captures perceptual sensitivity while controlling for response bias by incorporating both hit and false alarm rates (Green et al., 1966; Hautus et al., 2021; Stanislaw & Todorov, 1999). Thus, we chose *d′* scores over raw accuracy because it provides a more robust and unbiased estimate of participants’ ability to detect subtle phonetic changes, making it particularly well-suited for the two-alternative forced choice paradigm used in our study. *d’* scores were computed with the *psyphy* package (Knoblauch, 2023) run through R studio (R Core Team, 2023), using a differencing approach (Hautus et al., 2021). For Experiment 1, we additionally computed accuracy to compare potential ceiling or floor effects as a function of the SNR. We then used Jamovi (Doğan Şahin & Can Aybek, 2020) to ran linear mixed effect models (LMMs) as follow. For experiment 1, separately for *d’* and RTs we ran two LMMs (one for Exp. 1a with dialogues only and one for Exp. 1b with monologues only) with 2 factors [Semantic context (Intact / Scrambled) x SNR (-4 dB / - 6 dB)], baseline as covariate and subjects as random factor. For experiment 2, separately for *d’* and RTs we ran one LMM with 3 factors [Experimental design (a / b / c) x Semantic context (Intact / Scrambled) x Social context (Dialogue / Monologue)], baseline as covariate, and subjects as random factors. Because d′ scores and RTs were calculated per condition for each participant, the analysis was conducted on aggregated data at the subject level and item-level variability was not modeled. For each of these models, baseline performance was included as a covariate to take into account individual variability in performing the speech-in-noise task prior to manipulating the context. To evaluate the effects of the experimental factors, we extracted the estimated marginal means from these models and submitted them to fixed-effects omnibus tests in Jamovi. This approach allowed us to evaluate overall factor-level effects rather than contrasts against a specific reference level. In addition, we also ran the same LMM analysis for experiment 2 without including the baseline as a covariate, and we found that the model fit was better with the baseline than without the baseline (supplementary table I), although with very similar results (supplementary table II).

### 2. AQ analysis

In addition, we examined whether individual differences in processing conversations were related to autistic traits, as measured by the AQ. Specifically, we conducted one-tailed partial Spearman correlations, separately for dialogues and monologues, between individual AQ scores and the difference in performance between intact and scrambled conversations, while controlling for baseline performance as a potential confound. Here, we used a one-tail test because we had a clear hypothesis regarding the direction of our expected effect, a negative correlation, that would show that the lower the AQ score, the higher the difference of performance between intact and scrambled conversations. This hypothesis was directly driven from the results of an analogous study in the visual modality (Abassi & Papeo, 2022), that showed a negative correlation between individual AQ score and a measure of behavioral performances in visual processing of facing body dyads (as if interacting) as compared to non-facing body dyads, as well as other studies that showed similar relationship between AQ and visual sensitivity to other type of social stimuli (van Boxtel et al., 2017; Wyer et al., 2012; Yang et al., 2024). We then compared the difference of magnitudes of the Spearman *rhos* (Lenhard & Lenhard, 2014) between dialogues and monologues.

## III. RESULTS

### A. Experiment 1

Experiment 1 was conducted to choose a correct SNR for measuring our effects of interest in the subsequent experiments. To do so, based on prior in-the-lab pilots (unpublished data) we selected two SNRs of -6 and -4 dB. For the SNR of -6 dB we found a mean baseline accuracy of 61.9 % (+/- 1.9 Standard Error of the Mean [SEM]) for Exp. 1a and 64.5 % (+/- 1.8 SEM) for Exp. 1b, while for the SNR of -4 dB, we found a mean baseline accuracy of 73.2 % (+/- 1.8 SEM) for Exp. 1a and 72.3 % (+/- 1.8 SEM) for Exp. 1b. For the main task, for the SNR of -6 dB we found a mean average accuracy of 61.7 % (+/- 1.8 SEM) for Exp. 1a and 60.2 % (+/- 1.9 SEM) for Exp. 1b, while for the SNR of -4 dB we found a mean average accuracy of 78.1 % (+/- 1.8 SEM) for Exp. 1a and 77.1 % (+/- 1.5 SEM) for Exp. 1b.

Using *d’* scores, for Exp. 1a (dialogues only), the 2 SNR x 2 Semantic context LMM (fig. 2a) showed a main effect of SNR and no main effect of semantic context. Critically, it did show a significant interaction between SNR and Semantic context (table IIa). This interaction arises from a larger difference for intact vs. scrambled conversations for a SNR of -4 dB as compared to a SNR of -6 dB. Post-hoc *t*-tests showed that this difference was significantly higher for intact than scrambled conversations for a SNR of -4 dB (*t*(38) = 2.49, *p* = 0.017), while there was no difference for a SNR of -6 dB (*t*(38) = 0.35, *p* > 0.250). Using RTs (fig. 2b), we didn’t find any significant effects (table IIb). For Exp. 1b (monologues), we didn’t find any effects using d’ (table IIc; fig. 2c) or RTs (table IId; fig. 2d).

**Table II.**
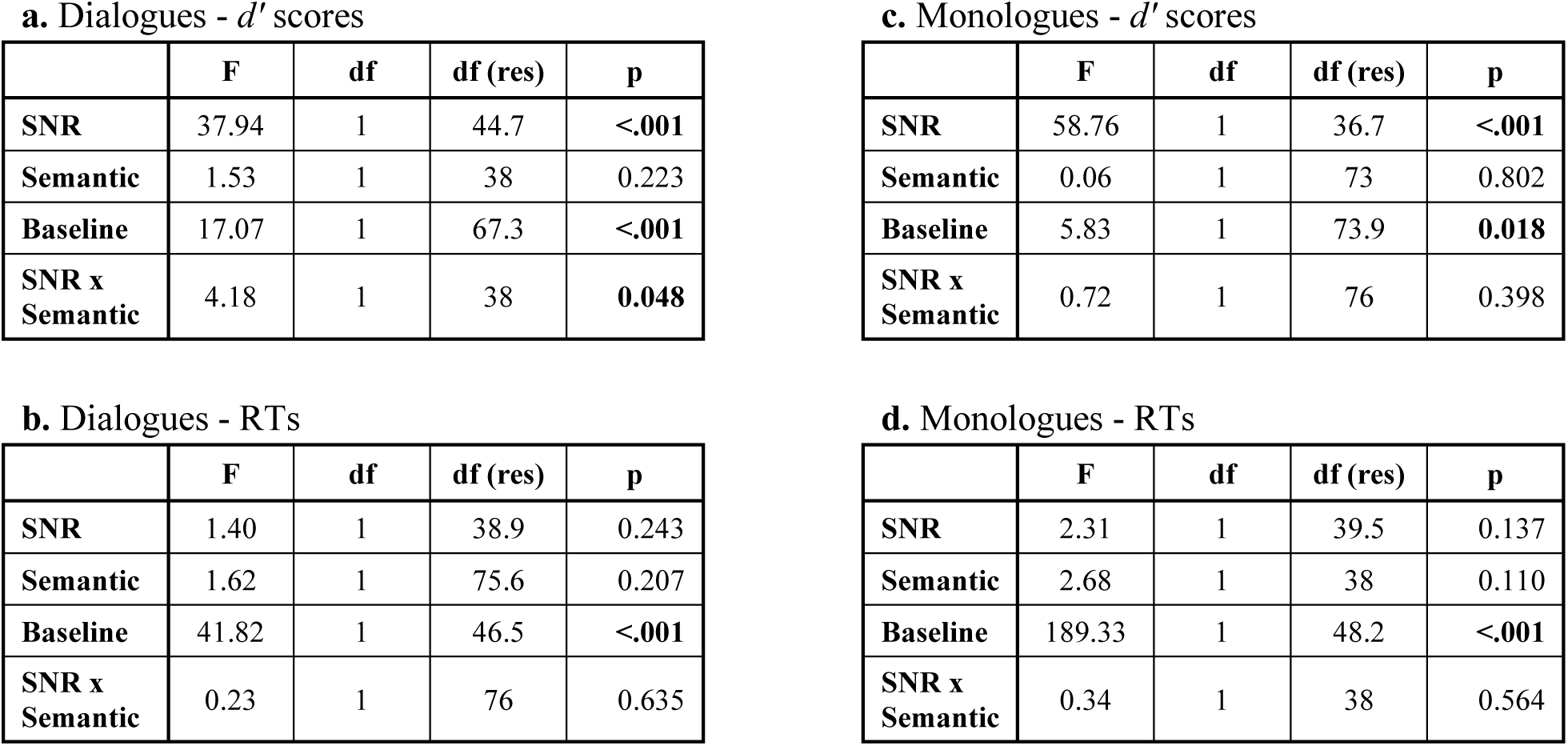
Results of the Linear Mixed Effect models with *d’* scores and RTs for dialogues (Exp. 1a: a.; b.) and monologues (Exp. 1b: c.; d.).

**a. Dialogues - $d'$ scores**
|  | F | df | df (res) | p |
| --- | --- | --- | --- | --- |
| SNR | 37.94 | 1 | 44.7 | <.001 |
| Semantic | 1.53 | 1 | 38 | 0.223 |
| Baseline | 17.07 | 1 | 67.3 | <.001 |
| SNR x Semantic | 4.18 | 1 | 38 | 0.048 |

**c. Monologues - $d'$ scores**
|  | F | df | df (res) | p |
| --- | --- | --- | --- | --- |
| SNR | 58.76 | 1 | 36.7 | <.001 |
| Semantic | 0.06 | 1 | 73 | 0.802 |
| Baseline | 5.83 | 1 | 73.9 | 0.018 |
| SNR x Semantic | 0.72 | 1 | 76 | 0.398 |

**b. Dialogues - RTs**
|  | F | df | df (res) | p |
| --- | --- | --- | --- | --- |
| SNR | 1.40 | 1 | 38.9 | 0.243 |
| Semantic | 1.62 | 1 | 75.6 | 0.207 |
| Baseline | 41.82 | 1 | 46.5 | <.001 |
| SNR x Semantic | 0.23 | 1 | 76 | 0.635 |

|  | F | df | df (res) | p |
| --- | --- | --- | --- | --- |
| SNR | 2.31 | 1 | 39.5 | 0.137 |
| Semantic | 2.68 | 1 | 38 | 0.110 |
| Baseline | 189.33 | 1 | 48.2 | <.001 |
| SNR x Semantic | 0.34 | 1 | 38 | 0.564 |

**Figure 2.**
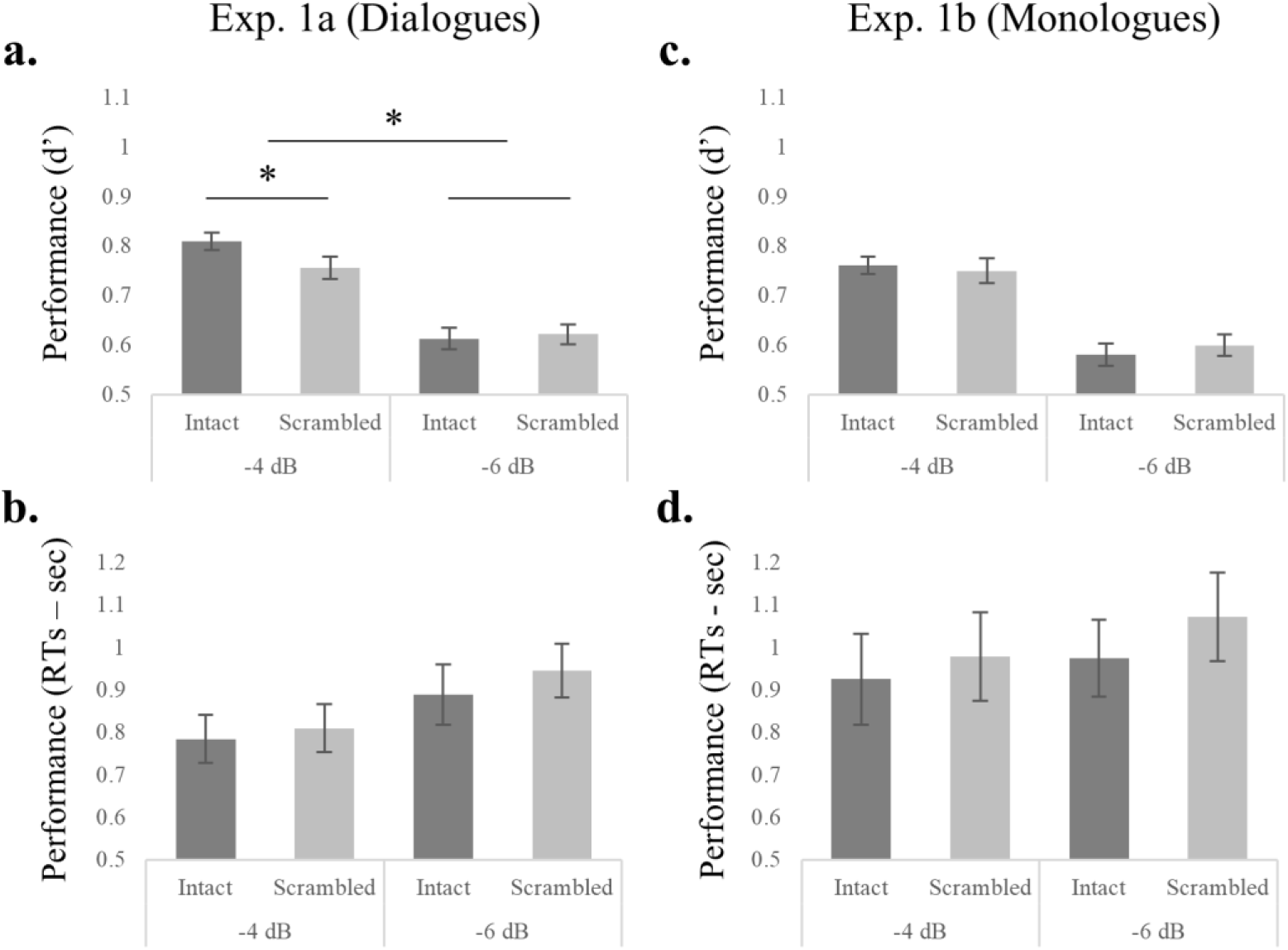
Behavioral data for experiment 1. Performances for intact and sentence-scrambled conversations using d-primes *(d’*) and response times (RTs) with SNRs of -4 dB and -6 dB for Exp. 1a (Dialogues only: **a.**; **b.**) and Exp. 1b (Monologues only: **c.**; **d.**). Error bars represent standard errors of the mean. *p ≤ 0.05.

To summarize, we first found that an SNR of -6 dB was producing accuracies in the range of 60-65 %, while an SNR of -4 dB was producing accuracies in the range of 72-78 %, that was a first indication that an SNR of -6 dB might be too close to chance level as compared to -4 dB. In addition, only with a SNR of -4 dB we found our effect of interest using *d’* (i.e., a significantly better performance for intact vs. scrambled dialogues in Exp. 1a). For this reason, we used an SNR of -4 dB for all subsequent experiments.

### B. Experiment 2

Raw accuracy scores for each condition were as follows: Intact Dialogues: 75.6 % (SD: 14.5); Scrambled Dialogues: 74.0 % (SD: 14.4); Intact Monologues: 75.2 % (SD: 14.2); Scrambled Monologues: 72.4 % (SD: 15.7); Baseline: 69.3 % (SD: 10.5). These values are reported for descriptive purposes only, all inferential statistics for performance were conducted on *d′* scores, which better reflect perceptual sensitivity in this task. Using *d’* scores (fig. 3a; Table IIIa), the 3 Design x 2 Semantic context x 2 Social context LMM showed only a significant main effect of semantic context, characterized by higher d-primes for intact than sentence scrambled conversations, and no interactions. Using RTs (fig. 3b; Table IIIb) the same analysis showed a significant effect of social context, characterized by lower RTs for dialogues than monologues, as well as a significant effect of semantic context, characterized by lower RTs for intact than sentence-scrambled conversations. It also showed a marginal trend for an interaction between experimental design and social factor, and no other effects or interactions.

**Table III.**
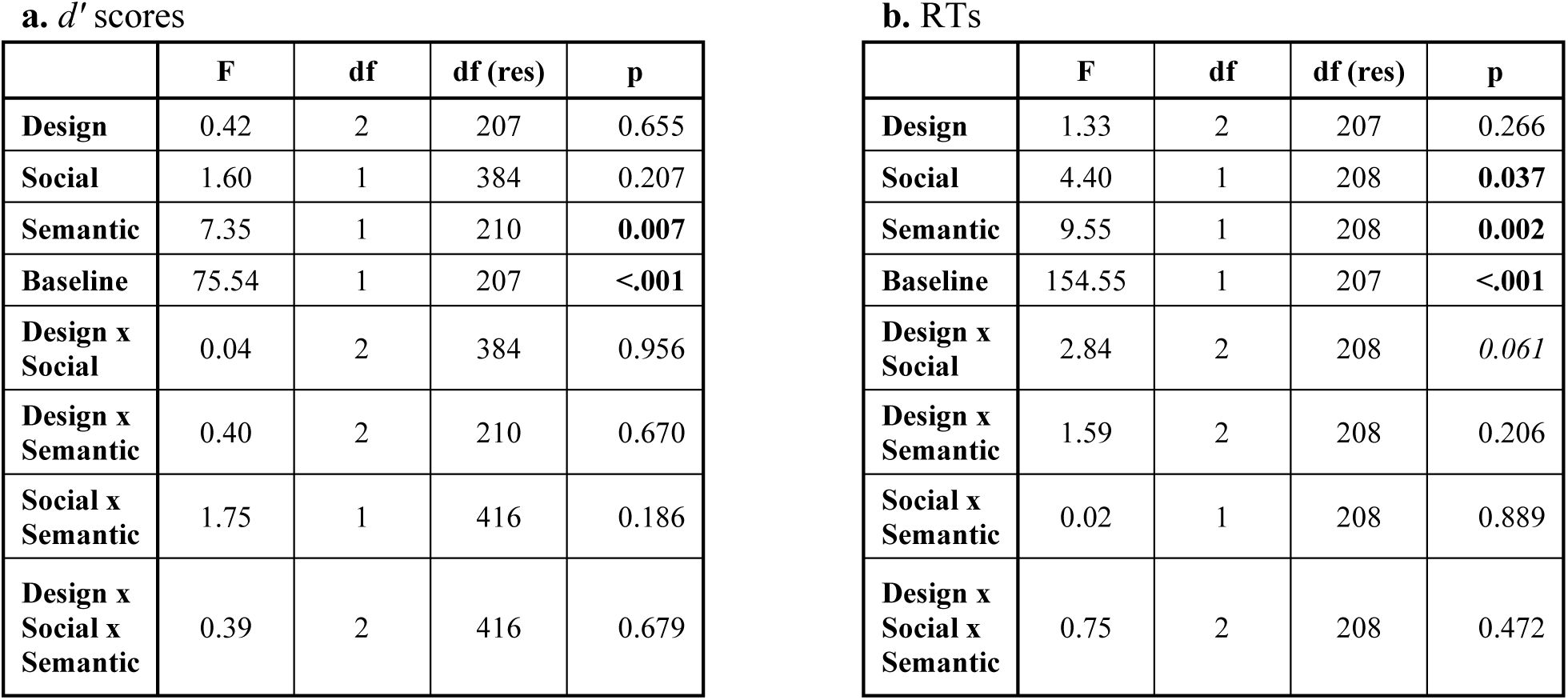
Results of the Linear Mixed Effect models with *d’* scores (a.) and RTs (b.) for Experiment 2.

**a. $d'$ scores**
|  | F | df | df (res) | p |
| --- | --- | --- | --- | --- |
| **Design** | 0.42 | 2 | 207 | 0.655 |
| **Social** | 1.60 | 1 | 384 | 0.207 |
| **Semantic** | 7.35 | 1 | 210 | **0.007** |
| **Baseline** | 75.54 | 1 | 207 | **<.001** |
| **Design x Social** | 0.04 | 2 | 384 | 0.956 |
| **Design x Semantic** | 0.40 | 2 | 210 | 0.670 |
| **Social x Semantic** | 1.75 | 1 | 416 | 0.186 |
| **Design x Social x Semantic** | 0.39 | 2 | 416 | 0.679 |

|  | F | df | df (res) | p |
| --- | --- | --- | --- | --- |
| **Design** | 1.33 | 2 | 207 | 0.266 |
| **Social** | 4.40 | 1 | 208 | **0.037** |
| **Semantic** | 9.55 | 1 | 208 | **0.002** |
| **Baseline** | 154.55 | 1 | 207 | **<.001** |
| **Design x Social** | 2.84 | 2 | 208 | 0.061 |
| **Design x Semantic** | 1.59 | 2 | 208 | 0.206 |
| **Social x Semantic** | 0.02 | 1 | 208 | 0.889 |
| **Design x Social x Semantic** | 0.75 | 2 | 208 | 0.472 |

**Figure 3.**
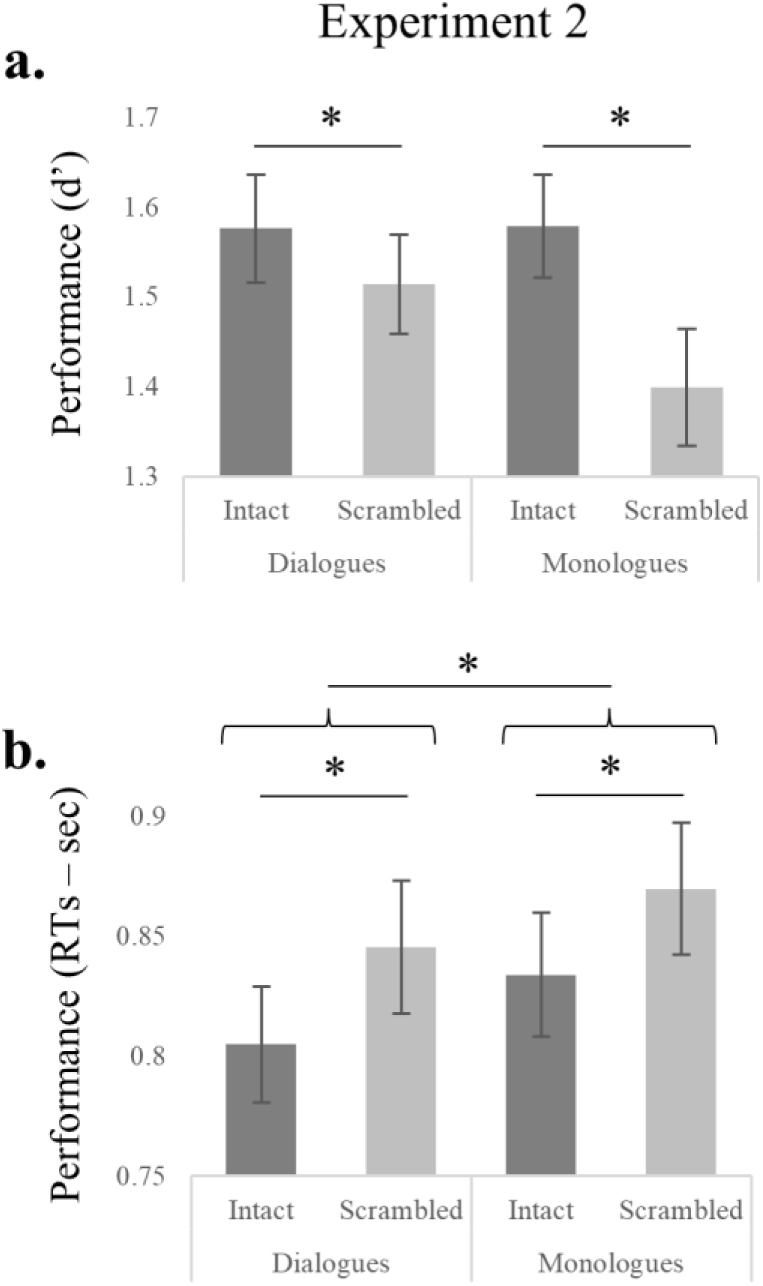
Behavioral data for experiment 2. Performances for intact and sentence-scrambled dialogues and monologues using d-primes (**a.**) and response times (**b.**). Error bars represent standard errors of the mean. Asterisks (*) represent main effects of the Omnibus test derived from our linear mixed models.

### C. Relationship between individual AQ scores and phonetic encoding

Using the data of experiment 2, we next examined whether individual differences in autistic traits (measured via AQ scores) were associated with sensitivity to semantic context, and whether this relationship varied depending on the social context of the conversation. To do so, we computed a difference score (Intact – Scrambled) reflecting the perceptual benefit of semantic coherence for each participant, separately for dialogues and monologues. These scores isolate the effect of semantic context within each social setting, allowing us to assess whether AQ modulates this effect differently depending on whether speech occurs in a socially interactive (dialogue) or non-interactive (monologue) context. Here, we didn’t find significant partial Spearman correlations using the *d’* score (Dialogues; *rho*(211) = -0.04; *p* > 0.250; Monologues; *rho*(211) = 0.07; *p* = 0.161) with no differences between dialogues and monologues (*z* = -1.1 *p* = 0.136). However, using RTs, only for dialogues and not for monologues we found that there was a trend for a negative correlation between AQ and processing of intact vs. scrambled conversations (Dialogues: rho(211) = -0.10, p = 0.067; Monologues: rho(211) = 0.02, p > 0.250). Furthermore, we also found a trend for a lower correlation for dialogues as compared to monologues (*z* = -1.38 *p* = 0.085).

## IV. DISCUSSION

We investigated how semantic and social contexts modulate phonetic encoding during processing of naturalistic conversations using a novel speech-in-noise task. First, we found that semantic context significantly enhanced phonetic encoding. Specifically, participants performed better (lower response time and better performance) when processing semantically coherent conversations compared to sentence scrambled ones. Second, social context also had a significant effect, with dialogues providing greater benefits (lower response time, but no effect on performance) to phonetic encoding than monologues. Finally, we found a trend for a negative correlation between individual differences in AQ traits and performances in processing intact vs. scrambled conversations for dialogues only.

Our finding that semantic coherence within a full conversation enhances phonetic encoding aligns well with existing research on predictive coding theories of auditory perception. Predictive coding posits that the brain generates top-down expectations based on semantic context to minimize mismatches between sensory input and prior knowledge (Clark, 2013; Friston, 2005). In our study, participants performed better when processing semantically coherent conversations, demonstrating the benefits of predictive mechanisms in noisy environments where bottom-up auditory signals are degraded. These results are consistent with previous work showing that semantic congruence facilitates words and phonemes identification while reducing cognitive load (Golestani et al., 2009, 2013; Sohoglu et al., 2012; Zekveld et al., 2011, 2012, 2013). However, our study extends these findings into a new domain by showing effects of semantic context at the larger scale of naturalistic, multi-sentence conversations. Here, while prior research has established the role of semantic context within short linguistic units (Broderick et al., 2019; Roland et al., 2012), our results suggest that predictive mechanisms also operate over longer timescales, integrating information across whole conversations. Indeed, in our paradigm, unlike most prior studies, the target sentence embedded in noise was not itself semantically anomalous; it was only anomalous or not with respect to the preceding conversational context.

While much of the prior research has framed these effects within the predictive coding framework, other complementary mechanisms may also explain the influence of semantic context on speech processing. First, semantic priming effects, whereby exposure to semantically related words activates shared neural networks, may also facilitate phonetic encoding by reinforcing linguistic expectations (Daltrozzo et al., 2011; Foss, 1982; Kutas & Federmeier, 2011). This process likely operates at multiple levels of representation, from lexical access to sentence-level coherence, allowing listeners to extract meaning even when speech is partially masked by noise. Secondly, semantic context may also interact with attentional mechanisms, prioritizing coherent information over incongruent or irrelevant input. Research on the cocktail party effect has shown that semantic predictability can help listeners segregate target speech from competing noise, even in multi-talker environments (Bronkhorst, 2015; Tóth et al., 2020). These findings suggest that semantic coherence not only aids phonetic encoding but also enhances the perceptual organization of auditory scenes, contributing to the broader goal of communication in complex environments. Interestingly, our study highlights the robustness of semantic effects across whole conversations, suggesting that listeners dynamically integrate information over longer timescales. This temporal dimension aligns well with research showing that discourse-level processing involves distinct neural mechanisms, including prefrontal and temporo-parietal networks, that support integration of meaning across sentences (Kutas & Federmeier, 2011; Menenti et al., 2012). Future research should explore how these long-range mechanisms interact with immediate predictive processes to shape phonetic encoding.

Our second and more novel findings pertain to social context effects, that showed better performances for dialogues over monologues. This result extends prior research emphasizing the role of interactional dynamics in speech processing. While the benefits of turn-taking and temporal predictability in dialogues are well-documented (Branigan et al., 2011; Fox Tree, 1999; Menenti et al., 2012; Pickering & Garrod, 2021; Tolins et al., 2018), this effect has been usually linked to the internal content of a dialogue as compared to a monologue, rather than to its intrinsic social value as an interaction. Indeed, dialogues usually contain information that monologues lack, such as discourses markers and/or different perspectives from multiple interlocutors, that may help in the understanding of the information that is shared (Pickering & Garrod, 2021). While this hypothesis can provide a partial explanation for this advantage in certain situations, it cannot apply to our study. Here, we used identical contents for both dialogues and monologues, with the only change being the vocal timbre of the speakers and their alternation or not. Furthermore, our task required the listener to identify if a word had been changed, but that word was not itself relevant to the content of the dialogue (e.g., see figure 1, in which the changed word was “true” vs “pure”: both of which fit the meaning of the sentence, and neither of which is related to the dialogue’s content). Thus, instead of emphasizing the content of the dialogue, we rather hypothesize that the presence of social cues, such as two speakers interacting with each other, triggers a sharpening of the acoustics of the sentences by the perceptual system, that is emphasizing some features of the auditory signal to facilitate comprehension, that would be higher for dialogues over monologues. This account suggests in turn that dialogues would provide additional cognitive engagement through social cues such as turn-taking and speaker interaction, which are absent in monologues.

Hence, dialogues, due to their interactive nature, would create a richer social context that may engage listeners more deeply by eliciting greater social attention and enhancing speech processing. This process could be mediated by neural systems specialized for processing socially relevant stimuli, such as the superior temporal sulcus (STS) and medial prefrontal cortex (Schilbach et al., 2006, 2008). This hypothesis is in line with recent findings in the domain of visual perception showing that a human body is better perceived when facing another body (i.e., mimicking a social interaction), as compared to facing in an opposite direction (Abassi et al., 2020; Papeo et al., 2017; Papeo & Abassi, 2019; Vestner et al., 2019, 2021), suggesting a domain-general effect that would enhance perceptual encoding in the presence of socially relevant settings, regardless of sensory modality.

Another unexplored factor is the role of joint attention in dialogues, where listeners anticipate and align with the speaker’s focus of attention. Studies in developmental psychology have shown that joint attention is critical for language acquisition and auditory processing (Mundy, 2018; Tomasello, 2005), suggesting that similar mechanisms may also operate in adult conversations to enhance comprehension. Additionally, the contingency of dialogue responses, where listeners predict speaker turns based on context, may activate reward systems (McNaughton et al., 2023; Stivers et al., 2009), further enhancing attention and processing efficiency. Interestingly, the absence of familiar speakers in our study highlights that interactional dynamics alone, independent of familiarity (Domingo et al., 2020; Johnsrude et al., 2013), can drive social context effects. Here, research on speaker familiarity would suggest that combining familiarity with dialogue-based cues could even amplify these benefits (Fleming et al., 2014; Yu et al., 2023). Future studies could investigate how these factors interact to optimize phonetic encoding in real-world scenarios.

An additional consideration is how our task relates to the broader concepts of intelligibility or phonetic encoding. While intelligibility is commonly defined as the successful recognition of spoken words under varying conditions (Mattys et al., 2012), phonetic encoding refers to the fidelity with which acoustic-phonetic information is mapped onto linguistic representations (Yi et al., 2019). These processes are closely linked: phonetic encoding is a foundational step that supports intelligibility. However, our task was designed to emphasize the former by focusing on subtle phonetic contrasts (e.g., “true” vs. “pure”, or “time” vs. “year”) where the alternative words were both grammatically plausible and semantically neutral with respect to the broader conversational context. In this way, listeners could not rely on sentence meaning to detect changes but instead had to encode and retain fine-grained phonetic detail. Thus, although our measure involves elements of intelligibility, particularly in noise, it targets how well listeners encode and retrieve phonetic forms under varying contextual conditions. By framing our results in terms of phonetic encoding, we highlight the influence of high-level semantic and social cues on early perceptual stages of speech processing.

One more point to highlight is that effects of social context were found only with response time, while effects of semantic context were found with both response time and accuracy. A first reason for this difference across tasks may be that our social condition task could be more fine-tuned (e.g., by using an adaptive SNR; see Soleymani et al., 2019) to better take into account individual variabilities that would be reflected in performance on the task and especially measures of accuracy (such as *d’* scores), that are more sensitive to floor or ceiling effects than RTs (Magnus et al., 2017; Willoughby et al., 2023). Another reason for the discrepancy could be due to the nature of the measured effects themselves. Indeed, it has been proposed that RTs and measures of accuracy may be affected by different attentional cues via distinct cognitive and/or neural mechanisms (Prinzmetal et al., 2005; van Ede et al., 2012 but see Mulder & Van Maanen, 2013). Thus, effects of social and semantic contexts may rely on different attentional cues, that may be reflected in measuring RTs and d-primes, while the exact nature of these different cues, if they exist, would need a more thorough exploration. On a last point, we note that we found only a marginal trend for an interaction of our conditions with the experimental design, showing that there is only a minimal effect of presentation order in our experiment.

We also note the lack of interaction between semantic and social contexts. Indeed, while both factors independently enhanced phonetic encoding, their effects appeared to operate additively rather than synergistically. This absence of interaction challenges theoretical frameworks suggesting that social and semantic cues are integrated to optimize speech processing (Menenti et al., 2012; Pickering & Garrod, 2021). One potential explanation lies in the modularity of processing streams for semantic and social information. Neural evidence suggests that semantic processing primarily engages regions such as the angular gyrus and inferior frontal gyrus (Humphries et al., 2007), while social interaction cues involve the STS and medial prefrontal cortex (Hasson et al., 2012; Schilbach et al., 2008). Although these networks are interconnected, their relative independence may explain why semantic and social effects did not interact in our study. It is possible that these processing streams converge at later stages, such as during integration of discourse-level meaning or higher-order inference-making, which were not directly tested in our task. Another possibility is that our task design, which isolated semantic and social factors, may have limited opportunities for these effects to interact. For example, the lack of speaker familiarity or shared conversational goals in our stimuli may have reduced the potential for synergistic effects. Indeed, research on common ground, the shared knowledge between conversational partners, has shown that mutual understanding enhances speech comprehension (Holler & Stevens, 2007; Horton & Keysar, 1996; Keysar et al., 2000). Future studies could test whether incorporating shared goals or familiar speakers facilitates interaction between semantic and social contexts. Finally, the additive effects observed in our study may reflect the distinct timescales at which semantic and social factors operate. Semantic coherence influences phonetic encoding already at the level of immediate word or sentence processing, while social context effects unfold over longer conversational exchanges. Understanding how these factors converge may require further investigation, particularly using experimental designs that manipulate both immediate and extended contextual cues, or other experimental methods such as neuroimaging.

An additional contribution of this study is the exploratory analysis of individual differences in social traits, measured by the AQ. While the negative correlation between AQ scores and the benefit of semantic coherence in dialogues over monologues was only a trend, and of low magnitude, it is consistent with previous findings suggesting that individuals with higher AQ scores may be less sensitive to social interaction cues (Bayliss et al., 2005). Similar negative associations between AQ and sensitivity to social stimuli have been reported in the visual modality (Abassi & Papeo, 2022; van Boxtel et al., 2017; Wyer et al., 2012; Yang et al., 2024), raising the possibility of a domain-general mechanism affecting how social cues are integrated across sensory systems. These observations do not contradict prior work showing enhanced attention to fine phonetic detail in individuals with higher AQ (Ota et al., 2015; Stewart & Ota, 2008; Yu, 2010), but may reflect a trade-off between local auditory precision and reduced sensitivity to social dynamics. While the present results provide only tentative support for this idea, they underscore the relevance of individual variability in social traits when investigating the interplay between social and semantic context in speech perception.

In sum, our findings contribute to several theoretical frameworks in auditory cognition and social perception. First, our results support predictive coding theories, demonstrating that both semantic and social contexts facilitate phonetic encoding, likely through top-down mechanisms. The robustness of these effects across large-scale conversations suggests that predictive mechanisms operate hierarchically, integrating information at multiple levels of linguistic and social processing. Second, our findings highlight the role of social interactional dynamics in engaging cognitive systems specialized for processing socially relevant stimuli, consistent with theories of dialogue-based processing and to neurocognitive models emphasizing the role of specialized neural circuits to social cognition (Pitcher & Ungerleider, 2021). Finally, the trend-level relationship between AQ and sensitivity to social context suggests a balance between enhanced detail-focused processing and reduced integration of social information, consistent with the weak central coherence hypothesis of autism, which proposes a cognitive style favoring local detail over global context (Happé & Frith, 2006; Hoy et al., 2004; Jolliffe & Baron-Cohen, 1999). However, the variability observed within AQ-related effects here should be understood in the context of a neurotypical population, since we drew our sample from a general population, and none of our participants had a known diagnosis of autism.

These findings also have practical implications for designing interventions and assistive technologies for individuals with hearing losses. For example, auditory training programs could incorporate semantically coherent narratives and simulate conversational dynamics to enhance speech processing in noisy environments. Similarly, hearing aids and other assistive devices could incorporate algorithms that prioritize interactional dynamics, such as turn-taking, to improve real-world communication outcomes. In a future research, we will extend the examination of the neural mechanisms underlying these effects using neuroimaging techniques to explore how different contexts modulate brain activity (see Landsiedel & Koldewyn, 2023; Olson et al., 2023). Here we predict that, using fMRI, we may find modulation and sharpening of brain activity in voice-selective regions of the auditory cortex, such as the STS, as a function of the semantic and/or social context. On another aspect, expanding the scope of this research to include older adults, individuals with hearing impairments, or non-native speakers could provide a broader understanding of how contextual factors impact diverse populations. Further explorations could also replicate our current findings using real human voices from naturalistic conversations, that might carry additional features of canonical social interactions. Finally, integrating cross-modal research on auditory and visual sensitivity for social interactions (see Abassi et al., 2024 for a review on social vision; see also Abassi & Papeo, 2024 for cross category research) could offer a more comprehensive perspective on disentangling the domain-general and domain-specific mechanisms underlying social cognition.

## V. SUPPLEMENTARY MATERIAL

See supplementary material at [URL will be inserted by AIP] for the stimuli set, additional figure and tables.

## ACKNOWLEDGMENTS

This research was supported by the Fondation Pour l’Audition through a postdoctoral fellowship to EA (FPA RD-2022-1) and by the Grand Prix Scientifique from the Fondation Pour l’Audition to RJZ (2021). RJZ holds a Canada Research Chair and is funded by the Canadian Institutes of Health Research..

## VI. AUTHOR DECLARATIONS

All authors declare that they have no financial or non-financial conflict of interests to disclose. They also declare that all procedures used in the current study were performed in compliance with the code of ethics of the World Medical Association’s Declaration of Helsinki and have been approved by the Faculty of Medicine Institutional Review Board of McGill University on the 3^rd^ of October 2022 (reference number: A08-B75-22B / 22-07- 116).

## VII. DATA AVAILIBILITY

**VIII.** The entire set of stimuli and materials for generation can be found online at www.zlab.mcgill.ca/downloads/Stimuli_from_Abassi_2025. A sample is also provided as supplementary files (*SuppPub2.zip*) and can be used under the term of a CC-BY-NC-SA 4.0 license. The collected online data are freely available upon email request to E.A.

## SUPPLEMENTARY MATERIALS

### Supplementary Information 1: Generation of multi-talker babble noise

#### Noise features

The features of the multi-talker babble noise was set as follow: 1) We used a total of 8 competing voices (4 females and 4 males), that is a good compromise between creating sufficient interference with the target voices while keeping a reasonable amount of competing voices (Cooke, 2006; Li & Lutman, 2006; Miller, 1947; Simpson & Cooke, 2005); 2) We selected the f0 of the competing voices so they would not overlap with the target voices, as having too close f0 between target and competing voices may interfere too much with the task (Assmann, 1999; Bird, 1997; Brokx & Nooteboom, 1982; David et al., 2017; Lee & Humes, 2012); 3) We selected a f0 range between ∼120 Hz and ∼203 Hz by steps of around one semi-tone between each voice (see supplementary Figure 1), so the range would be large enough to cover a representative range of f0 in the general population (see Cristina Oliveira et al., 2021).

#### Noise distractor sentences generation

To generate the content of the noise distractor sentences, we used ChatGPT to generate 24 semantically anomalous sentences in English. Anomalous sentences were used here to eliminate the possibility that participants might extract an entire meaningful sentence from a speaker other than the target while keeping correct grammar (Van Engen & Bradlow, 2007). An example of the ChatGPT prompts used is *“Write X semantically anomalous but syntactically correct sentences”* and an example of the anomalous sentences generated is *“Her purple umbrella whistled a dolphin”.* We then selected eight voices from the list proposed by Google text-to-speech AI: 4 females voices (*’en-US-News-K’; ‘en-US-News-L’; ‘en-US-Wavenet-E’; ‘en-US-Wavenet-C’*) and 4 male voices (*’en-US-Wavenet-D’; ‘en-US-News-N’; ‘en-US-Wavenet-I’; ‘en-US-News-M’*). Prior to generating the audio speech for the noise, we set the f0 parameter for each voice as follows. For all voices we used the vowel /ɑ/ as a reference (Cristina Oliveira et al., 2021; De Krom, 1994), then for each voice we measured the original f0 of the voice for the duration of this vowel using the built-in function ‘pitch’ of Matlab (R2022b; The MathWorks Inc.) and time windows of 52 ms., and we defined the f0 as the median of the f0 measured for all time windows. Using Matlab, we then calculated by how much we should increase or decrease the original f0 of the voice to get to the custom f0 we wanted.

### Supplementary Information 2: Check of optimal on-line experimental conditions

Prior to beginning an experiment, participants went through a series of checks to ensure optimal ‘auditory hygiene’ while performing the tasks. To do so, we used guidelines and scripts from (Zhao et al., 2022). More specifically, participants first heard multi-talker babble noise alone without target sentences and they were instructed to adjust their audio volume to a comfortable level, so they would not be exposed to overly loud sounds during the experiment. They then performed a headphone screening using Antiphase tone (Woods et al., 2017) to confirm that they were wearing headphones. More specifically, for this task participants had to correctly detect an acoustic target in an intensity-discrimination task. On each trial, they were presented with three consecutive 200 Hz sinusoidal tones in random order, and they were asked to determine which of the tones was perceptually the softest. Two of the tones were presented diotically: 1) the ‘standard’, and 2) the ‘target’ that was presented at -6 dB relative to the standard. The third tone (the ‘foil’) had the same amplitude as the standard but was presented dichotically such that the left and right signals had opposite polarity (anti-phase, 180°). Over headphones, the standard and foil should have the same loudness, making the target clearly distinguishable as softer. In contrast, over loudspeakers the left and right signals interact destructively before reaching the listener’s ears, resulting in a weaker acoustic signal at both ears, and thus a weaker loudness sensation for the foil than the target, causing participants to respond incorrectly. If participants failed 5 out of 6 trials, they were proposed to redo the headphone check a second time. If they failed the headphone check a second time, they were not allowed to carry on with the experiment and the study automatically stopped for them.

---

**Supplementary Figure 1.**
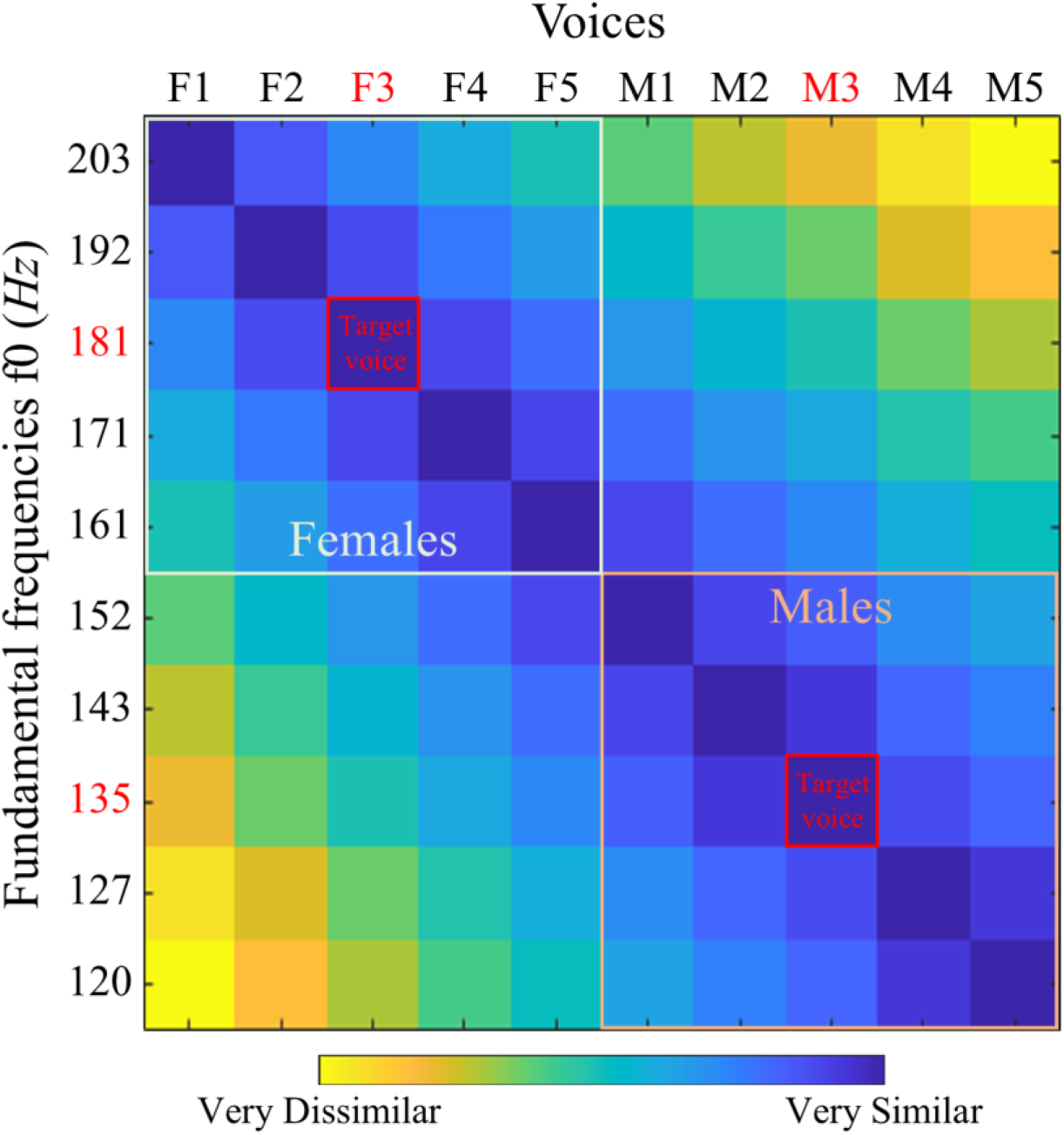
Dissimilarity matrix for Euclidean distances between fundamental frequencies of the voices. Captions in red represent the target voices for female (F3) and male (M3). Captions in black represent the concurrent voices of females (F1, F2, F4, F5) and males (M1, M2, M4, M5) that are used in the multi-talker babble noise

**Supplementary Table I.**
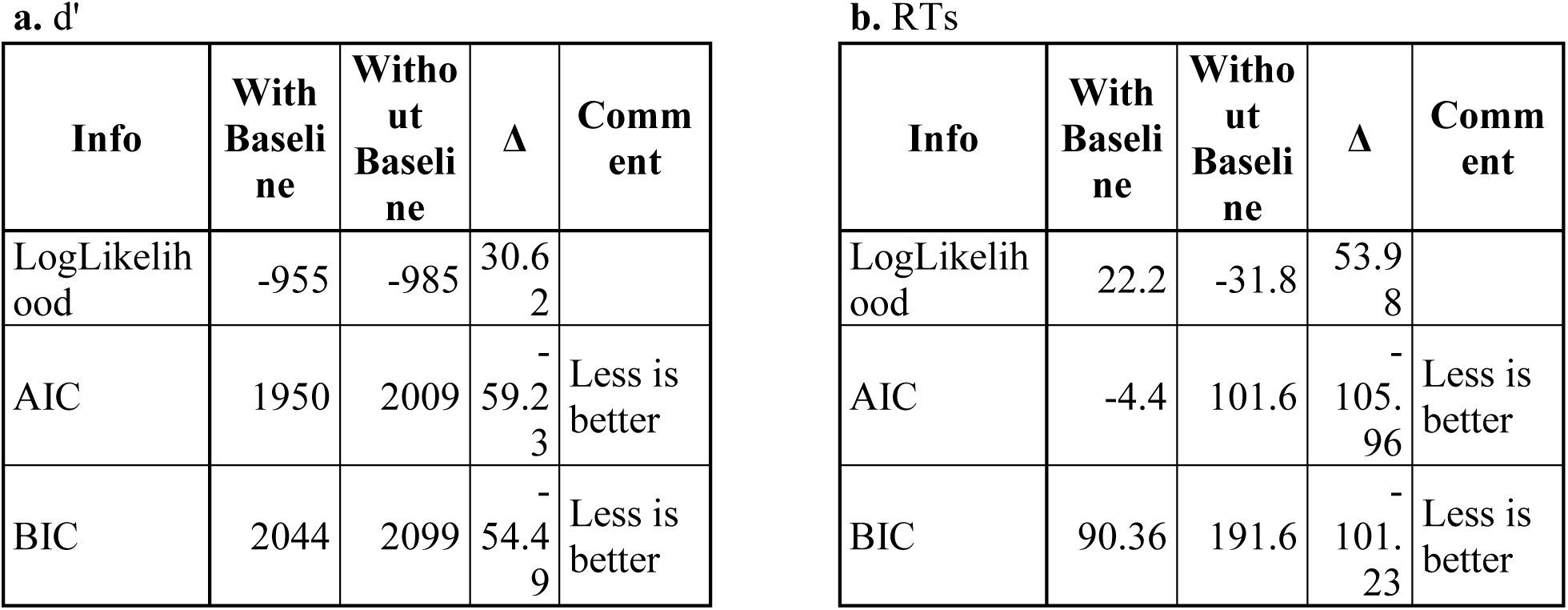
Comparison of Linear Mixed Effect models fits with and without including the baseline as a covariate.

**Supplementary Table II.**
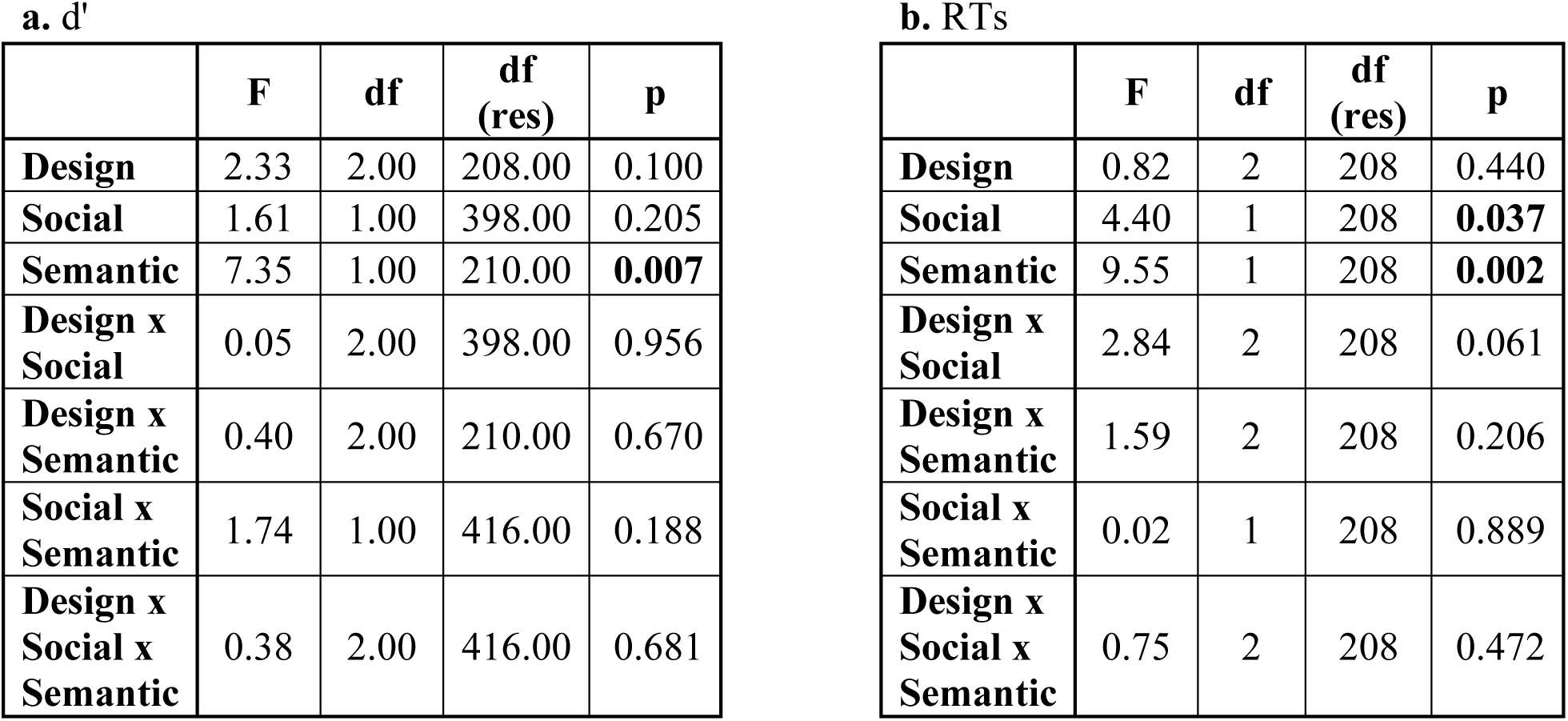
Results of the Linear Mixed Effect models with d’ (a.) and RTs (b.) for Experiment 2 without baseline as a covariate.

**a. d'**
|  | F | df | df (res) | p |
| --- | --- | --- | --- | --- |
| **Design** | 2.33 | 2.00 | 208.00 | 0.100 |
| **Social** | 1.61 | 1.00 | 398.00 | 0.205 |
| **Semantic** | 7.35 | 1.00 | 210.00 | **0.007** |
| **Design x Social** | 0.05 | 2.00 | 398.00 | 0.956 |
| **Design x Semantic** | 0.40 | 2.00 | 210.00 | 0.670 |
| **Social x Semantic** | 1.74 | 1.00 | 416.00 | 0.188 |
| **Design x Social x Semantic** | 0.38 | 2.00 | 416.00 | 0.681 |

**b. RTs**
|  | F | df | df (res) | p |
| --- | --- | --- | --- | --- |
| **Design** | 0.82 | 2 | 208 | 0.440 |
| **Social** | 4.40 | 1 | 208 | **0.037** |
| **Semantic** | 9.55 | 1 | 208 | **0.002** |
| **Design x Social** | 2.84 | 2 | 208 | 0.061 |
| **Design x Semantic** | 1.59 | 2 | 208 | 0.206 |
| **Social x Semantic** | 0.02 | 1 | 208 | 0.889 |
| **Design x Social x Semantic** | 0.75 | 2 | 208 | 0.472 |

